# Serotonin Enhances Neurogenesis Biomarkers, Hippocampal Volumes, and Cognitive Functions in Alzheimer’s Disease

**DOI:** 10.1101/2024.10.02.616371

**Authors:** Ali Azargoonjahromi, Alzheimer’s Disease Neuroimaging Initiative

**Author notes:** **Corresponding Author:** Ali Azargoonjahromi. Data used in preparation of this article were obtained from the Alzheimer’s Disease Neuroimaging Initiative (ADNI) database (adni.loni.usc.edu). As such, the investigators within the ADNI contributed to the design and implementation of ADNI and/or provided data but did not participate in analysis or writing of this report. A complete listing of ADNI investigators can be found at: http://adni.loni.usc.edu/wp-content/uploads/how_to_apply/ADNI_Acknowledgement_List.pdf.

## Abstract

Research on serotonin reveals a lack of consensus regarding its role in brain volume, especially concerning biomarkers linked to neurogenesis and neuroplasticity, such as ciliary neurotrophic factor (CNTF), fibroblast growth factor 4 (FGF-4), bone morphogenetic protein 6 (BMP-6), and matrix metalloproteinase-1 (MMP-1) in Alzheimer’s disease (AD). This study aimed to investigate the influence of serotonin on brain structure and hippocampal volumes in relation to cognitive functions in AD, as well as its link with biomarkers like CNTF, FGF-4, BMP-6, and MMP-1. Data from the ADNI included 133 AD participants. Cognitive function was assessed using CDR-SB, serotonin levels were measured with the Biocrates AbsoluteIDQ p180 kit and UPLC-MS/MS, and neurotrophic factors and biomarkers were quantified using multiplex targeted proteomics. Voxel-Based Morphometry (VBM) analyzed gray matter volume changes via MRI. Statistical analyses employed Pearson correlation and Bootstrap methods, with p-values < 0.05 or 0.01 considered significant. The analysis revealed a significant positive correlation between serotonin levels and total brain volume (r = 0.179, p = 0.039) and hippocampal volumes (right: r = 0.181, p = 0.037; left: r = 0.217, p = 0.012). Besides, higher serotonin levels were associated with improved cognitive function, evidenced by a negative correlation with CDR-SB scores (r = -0.198, p = 0.023). Furthermore, total brain volume and hippocampal volumes showed significant negative correlations with CDR-SB scores, indicating that greater cognitive impairment was associated with reduced brain volume (total: r = -0.223, p = 0.010; left: r = -0.246, p = 0.004; right: r = -0.308, p < 0.001). Finally, serotonin levels were positively correlated with BMP-6 (r = 0.173, p = 0.047), CNTF (r = 0.216, p = 0.013), FGF-4 (r = 0.176, p = 0.043), and MMP-1 (r = 0.202, p = 0.019), suggesting a link between serotonin and neurogenesis and neuroplasticity. In conclusion, increased serotonin levels are associated with improved cognitive function, increased brain volume, and elevated levels of neurotrophic factors and biomarkers—specifically CNTF, FGF-4, BMP-6, and MMP-1—that are related to neurogenesis and neuroplasticity in AD.

**Graphical Abstract:** 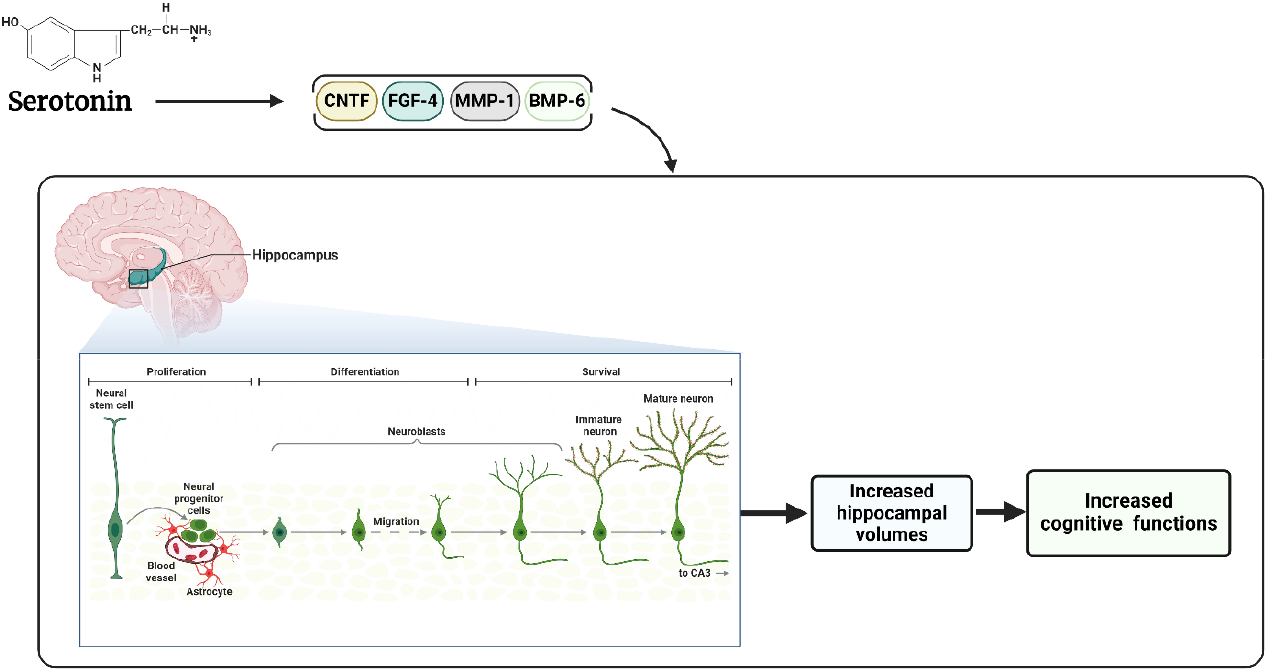

Serotonin plays a key role in cognitive function, with higher levels linked to increased brain and hippocampal volumes and better cognitive performance, as shown by lower CDR-SB scores (indicating less cognitive impairment). In addition, serotonin is positively associated with neurotrophic factors and biomarkers like ciliary neurotrophic factor (CNTF), fibroblast growth factor 4 (FGF-4), bone morphogenetic protein 6 (BMP-6), and matrix metalloproteinase-1 (MMP-1), which are involved in neurogenesis and neuroplasticity. These findings suggest that elevated serotonin levels contribute to brain health, cognitive function, and the promotion of neural growth and plasticity, particularly in Alzheimer’s disease (AD).

## 1. Introduction

Serotonin (5-hydroxytryptamine, 5-HT) is a vital neurotransmitter that regulates mood, promotes neurogenesis, and supports neuroplasticity, particularly in the hippocampus [1-4], which is essential for learning and emotional regulation [5, 6]. Known as the “feel-good” chemical, serotonin is predominantly found in the brain and gastrointestinal tract, with about 90% located in the gut. It is synthesized from the amino acid tryptophan and influences various physiological functions, including mood regulation, bowel movement control, blood vessel constriction, appetite, sleep, and cognition [7, 8]. Notably, serotonin’s role in regulating neurogenesis and synaptic plasticity makes it a crucial focus in research on mood disorders like depression [9], as well as neurocognitive diseases such as Alzheimer’s disease (AD) [10].

Research has shown that serotonin induces neurogenesis, the formation of new neurons, primarily in the dentate gyrus (DG) of the hippocampus and the subventricular zone (SVZ) [11-14]. Indeed, elevated serotonin levels, often achieved through antidepressants such as selective serotonin reuptake inhibitors (SSRIs), have been demonstrated to enhance the generation of new neurons in these areas [15, 16]. Accordingly, serotonin and its receptors, including 5-HT1A, 5-HT1B, and 5-HT2, are crucial for promoting the growth of neural progenitor cells, which ultimately leads to the formation of new neurons [1, 12, 13, 17-19]. For instance, the activation of the 5-HT1A receptor has been shown to enhance neuron production in both the DG and SVZ; likewise, the 5-HT2A and 5-HT2C receptors influenced neurogenesis in various brain regions, underscoring serotonin’s diverse roles in neuronal development [11].

Despite such findings, a study found that lowering central serotonin levels in adulthood, achieved by depleting serotonergic neurons or inactivating serotonin synthesis, enhances adult hippocampal neurogenesis [20]. Similarly, a surprising finding in a transgenic rat model (TetO-shTPH2) suggested that reduced serotonin levels may actually promote increased neurogenesis, potentially serving as a compensatory mechanism in response to stress or injury. This challenges the prevailing view that higher serotonin levels consistently enhance neurogenesis [21]. Thus, such complexities highlight the nuanced role of serotonin in regulating neurogenesis under various conditions, including stress and neurodegeneration.

Beyond neurogenesis, serotonin can enhance neuroplasticity—the brain’s ability to form new neural connections—by interacting with brain-derived neurotrophic factor (BDNF), which supports neuron growth and synapse formation [14, 22, 23]. Notably, this interaction strengthens synaptic connections in the hippocampus, promotes neurogenesis and stress resilience—as shown in both human and animal studies—and plays a crucial role in cognitive functions due to increased BDNF expression [23-26]. Noteworthy, regarding serotonin’s role in other neurotrophic and nerve growth factors such as ciliary neurotrophic factor (CNTF), fibroblast growth factor 4 (FGF-4), bone morphogenetic protein 6 (BMP-6), and matrix metalloproteinase-1 (MMP-1), there has been, to our knowledge, no research conducted to clarify the specific interactions and regulatory mechanisms between serotonin and these factors concerning neurogenesis, neuroplasticity, and metabolic regulation. Therefore, further studies are essential to understand how serotonin influences these neurotrophic factors and their potential implications for neurodevelopment and neurodegenerative disorders. This current study aimed to address this gap.

In AD, the serotonin system was found to influence on key pathological processes such as β-amyloid and tau protein aggregation. Evidence suggested that targeting serotonin receptors may not only improve cognitive function but also address the root causes of AD-related dementia [10]. However, s evidence from zebrafish models revealed that serotonin can negatively regulate BDNF expression [27], which contrasts with other studies showing that serotonin and BDNF work synergistically to enhance neuroplasticity and neurogenesis [22-26]. This discrepancy suggests that serotonin may behave differently in neurodegenerative diseases like AD compared to typical neuroplastic or antidepressant mechanisms. This study aimed to explore the interaction between serotonin levels and neurotrophic factors and biomarkers related to neurogenesis and neuroplasticity, which has yet to be elucidated.

The former studies underscored the critical role of serotonin in regulating brain structure and function, particularly gray matter volume (GMV) and hippocampal volume. A positive correlation existed between serotonin receptor binding, especially the 5-HT1A receptor, and GMV, indicating these receptors influence brain integrity [28]. Likewise, in psychiatric conditions, altered serotonin receptor densities are linked to decreased GMV and hippocampal volumes, impaired neurogenesis, and structural brain changes, thereby contributing to mood disturbances and cognitive deficits [29, 30]. Of note, genetic variants, such as His452Tyr in the 5-HT2A receptor, were linked to reduced GMV in the hippocampus and poorer memory performance, highlighting the genetic influence on serotonin’s effects [31]. However, the correlation between 5-HT1A receptor binding and GMV was absent in individuals with autism spectrum disorder [32]. Overall, these findings emphasize serotonin’s multifaceted role in brain structure and function, warranting further investigations.

This study aimed to investigate, to the best of our knowledge for the first time, how serotonin influences brain structure and hippocampal volumes, along with their connections to cognitive functions in AD.

Besides, it sought to explore the relationship between serotonin and neurotrophic factors and biomarkers, such as CNTF, FGF-4, BMP-6, and MMP-1, pertained to neurogenesis and neuroplasticity.

## 2. Methods and Materials

Data for this article were obtained from the Alzheimer’s Disease Neuroimaging Initiative (ADNI) database (www.adni.loni.usc.edu), launched in 2003 by Principal Investigator Michael W. Weiner, MD. ADNI aims to determine if MRI, PET, biological markers, and clinical assessments can track the progression of MCI and early AD. The study recruited 133 participants who met the ADNI criteria for the AD group.

### 2.1. Cognitive Assessment

Among the three cognitive assessments—CDR, ADAS-Cog, and MMSE—this study specifically utilized the CDR-SB to establish correlations with other factors. In contrast, both the MMSE and ADAS-Cog were employed to provide a broader analysis of the participants’ overall cognitive functions. This distinction allows for a nuanced understanding of cognitive impairments in the context of AD, highlighting the unique role of each assessment tool in the evaluation process.

The clinical dementia rating (CDR) [33] scale was used to assess the overall severity of dementia symptoms across multiple domains, including memory, orientation, judgment, and personal care. The evaluation process involved a semi-structured interview with the participant and an interview with an informant, typically a family member or caregiver. Based on these interviews, scales ranged from 0 (no impairment) to 3 (severe impairment). The ratings from each domain were then combined to produce a global CDR score, which classifies participants into categories ranging from normal cognition, to mild cognitive impairment or dementia [34]. The CDR-sum of boxes (CDR-SB) score ranges from 0 to 18, with higher scores indicating greater cognitive and functional impairment. A score of 0 reflects no impairment, while scores between 0.5 and 18 represent varying levels of mild to severe dementia [35].

The Alzheimer’s disease assessment scale-cognitive subscale (ADAS-Cog) [36, 37] was specifically designed to measure the severity of cognitive symptoms associated with AD. It includes a series of cognitive tests that assess various domains such as memory, language, and praxis (motor coordination). Participants were asked to perform tasks like word recall, object naming, and copying geometric figures. The total score on the ADAS-Cog can range from 0 to 70, with higher scores indicating more severe cognitive impairment. This longitudinal tracking was crucial for understanding how cognitive decline progresses in individuals with MCI or AD [37].

The mini-mental state examination (MMSE) [38] is a widely used screening tool for assessing cognitive function, primarily in older adults. It comprises 30 questions that evaluate various cognitive domains, including orientation, attention, memory, language, and visuospatial skills. The MMSE score ranges from 0 to 30, with higher scores indicating better cognitive function. A score of 24 or lower typically suggests cognitive impairment, with lower scores often correlating with greater severity of dementia. The MMSE is commonly used in clinical settings to help diagnose and monitor cognitive decline [39, 40].

### 2.2. Serotonin Measurement

The measurement of serotonin levels in human serum samples was carried out using the Biocrates AbsoluteIDQ p180 kit, which quantifies over 180 metabolites, including serotonin. The process began with sample preparation, where 10 µL of serum was added to each well of a 96-well plate containing an internal standard for accurate quantitation. The plate was then dried under nitrogen, followed by derivatization with phenyl isothiocyanate to label biogenic amines, including serotonin. The samples were eluted with 5 mM ammonium acetate in methanol for extraction and diluted with 40% methanol in water for UPLC analysis.

Serotonin was analyzed using UPLC-MS/MS, with separation achieved through a Waters Acquity UPLC system using a C18 column. A gradient of 0.1% formic acid in water and acetonitrile was used, and detection occurred in Multiple Reaction Monitoring (MRM) mode with a Xevo TQ-S triple quadrupole mass spectrometer. The transition from precursor to product ions was monitored, with stable isotope-labeled standards ensuring accuracy. Data processing was done using TargetLynx™ software, with final analysis in Biocrates MetIDQ™ software. Calibration standards and quality control (QC) measures, including Golden West Serum, NIST SRM-1950 plasma, and a Study Pool QC, were incorporated to maintain accuracy. The QC pool was injected before and after every batch to check for potential batch effects, ensuring reliable results.

### 2.3. Neurotrophic and Growth Factors Measurement in Plasma Samples

In the ADNI, the measurement of neurotrophic and growth factors in plasma involved stringent QC procedures. QC measures included spiking blank human plasma with extracts from cell cultures expressing individual analytes, with low, medium, and high QC levels established for nearly all analytes and run in duplicate on 15 plates. A total of 1,065 500 μL EDTA plasma samples were processed, collected from participants after overnight fasting, within 120 minutes to ensure sample integrity.

The laboratory employed a multiplex targeted proteomics approach for quantifying analytes, which allowed for simultaneous measurement of multiple proteins in a single sample. Samples were randomized for analysis at the RBM facility, which remained blind to the clinical data. Both baseline and one-year follow-up samples were included, targeting individuals with MCI and AD. QC results were carefully monitored, with analytes showing CV values above 25% flagged for caution. The methodology addressed potential discrepancies such as outliers and samples below the least detectable dose (LDD). Statistical analyses adhered to a predefined plan, highlighting the importance of consulting trained statisticians for accurate interpretation.

### 2.4. Neuroimaging Technique (Voxel-Based Morphometry (VBM))

Voxel-based morphometry (VBM) using MRI data is a technique applied for the quantitative analysis of brain structural differences. VBM focuses on detecting regional volume changes in gray matter by analyzing the voxel-wise comparison of brain anatomy across different subjects, particularly in the context of neurodegenerative diseases like AD [41].

VBM involves several critical preprocessing steps to ensure accurate and reliable comparison of brain structures across subjects. First, high-resolution T1-weighted MRI images undergo denoising and bias correction to reduce noise and correct intensity variations. The brain images are then segmented into gray matter, white matter, and cerebrospinal fluid (CSF) compartments using a Bayesian framework. After segmentation, spatial normalization aligns the images to a standardized space (typically MNI), allowing for comparability across subjects. Finally, the segmented gray matter images are smoothed with a Gaussian kernel, improving statistical power and accommodating anatomical variability. VBM analysis is then performed using voxel-wise comparisons across different groups (e.g., AD patients versus controls), often using statistical parametric mapping (SPM) software. Statistical analysis follows, adjusting for multiple comparisons to ensure the reliability of results. This standardized pipeline ensures consistent, large-scale analysis of brain structure in neurodegenerative diseases.

### 2.5. Statistical Analysis

All statistical analyses were performed using IBM SPSS Statistics version 27. Descriptive statistics, including means, standard deviations, and percentages, were calculated for the demographic characteristics and cognitive scores of the participants. Pearson correlation coefficients were computed to examine the relationships between serotonin levels and various brain volume measurements, including total brain volume and hippocampal volumes. The significance of these correlations was assessed using a two-tailed test, with a p-value of less than 0.05 or 0.01 considered statistically significant. Bootstrap analysis was utilized to evaluate the stability of estimates and to provide bias-corrected confidence intervals for the correlations. A 95% confidence interval was calculated to further assess the precision of the estimates. Covariance analyses were conducted to understand the degree to which changes in serotonin levels and cognitive function scores occurred in relation to each other. These analyses aimed to highlight the interrelationships between serotonin levels, brain volume, and cognitive function, reinforcing the study’s findings regarding the potential role of serotonin in neuroplasticity and cognitive health in AD.

## 3. Results

A total of 133 participants diagnosed with AD were included in the study, comprising 79 males and 54 females. Participants’ ages ranged from 56 to 90 years, with a mean age of 76.20 years (SD = 7.39). Cognitive function was measured using multiple tools: the CDR-SB (mean = 4.88, SD = 4.14), the MMSE (mean = 22.11, SD = 4.28), and the ADAS-Cog (mean = 17.02, SD = 7.22). (Table 1)

**Table 1:** Demographic characteristics of the participants.

| Variable | N | Mean (SD) | Min | Max |
| --- | --- | --- | --- | --- |
| **Age (Y/O)** | N/A | 76.20 (7.39) | 56 | 90 |
| **Handedness** | N/A | 1.08 (0.28) | 1 | 3 |
| - Right-handed | 123 | N/A | N/A | N/A |
| - Left-handed | 10 | N/A | N/A | N/A |
| **Gender** | N/A | 1.41 (0.49) | 1 | 2 |
| - Males | 79 | N/A | N/A | N/A |
| - Females | 54 | N/A | N/A | N/A |
| **Education** | N/A | 15.20 (3.04) | 8 | 20 |
| **CDR-SB** | N/A | 4.88 (4.14) | 0 | 18 |
| **MMSE** | N/A | 22.11 (4.28) | 5 | 30 |
| **ADAS-Cog** | N/A | 17.02 (7.22) | 3 | 43 |

### 3.1. Serotonin Level Correlated with Total Brain Volume and Bilateral Hippocampal Volumes

The findings revealed correlations between serotonin levels and various brain volume measurements, including total brain volume and left and right hippocampal VBM. There was a significant positive correlation between serotonin and total brain volume (r = 0.179, p = 0.039), as well as between serotonin and right hippocampal VBM (r = 0.181, p = 0.037). A moderate positive correlation was observed between serotonin and left hippocampal VBM (r = 0.217, p = 0.012), also statistically significant. Besides, total brain volume showed positive correlations with both left hippocampal VBM (r = 0.172, p = 0.048) and right hippocampal VBM (r = 0.203, p = 0.019). Notably, left and right hippocampal VBM are strongly correlated (r = 0.870, p < 0.001). The results, supported by bootstrap confidence intervals and standard errors, suggested positive relationships between serotonin and brain volume metrics, particularly in the hippocampal regions. (Table 2)

**Table 2:** Correlations between serotonin level and various brain volume measurements.

| Variables | Serotonin | Total Brain Volume | Left Hippo VBM | Right Hippo VBM |
| --- | --- | --- | --- | --- |
| **Serotonin** | 1 | **0.179\*** ($p = 0.039$) | **0.217\*** ($p = 0.012$) | **0.181\*** ($p = 0.037$) |
| **Total Brain Volume** | **0.179\*** | 1 | **0.172\*** ($p = 0.048$) | **0.203\*** ($p = 0.019$) |
| **Left Hippocampal VBM** | **0.217\*** | **0.172\*** | 1 | **0.870\*\*** ($p < 0.001$) |
| **Right Hippocampal VBM** | **0.181\*** | **0.203\*** | **0.870\*\*** | 1 |

\*Correlation is significant at the 0.05 level (2-tailed). \*\*Correlation is significant at the 0.01 level (2-tailed).

### 3.2. Serotonin Level Improved Cognitive Functions Measured by CDR-SB

The findings showed the correlation between serotonin levels and CDR-SB scores in 133 participants. The Pearson correlation coefficient of -0.198 indicated a negative relationship, meaning higher serotonin levels are associated with lower CDR-SB scores, which reflect greater cognitive functions. This correlation was statistically significant (p = 0.023), suggesting a less than 5% probability that the result was due to chance. The sum of squares for serotonin was 21.593, and for CDR-SB, it was -25.287, indicating their interrelatedness.

Covariance values indicated the degree to which two variables change together. A covariance of 0.164 for serotonin meant that as serotonin levels increase, there was a tendency for the values of the other variable (CDR-SB) to change positively. Conversely, a covariance of -0.192 for CDR-SB suggested that as CDR-SB scores increase (indicating more cognitive impairment), serotonin levels tended to decrease.

Bootstrap analysis provided insights into the stability of these estimates. A bias of 0 for serotonin indicated that the estimate of its relationship with CDR-SB was unbiased, while a bias of -0.004 for CDR-SB suggested a slight negative adjustment. Standard errors reflected the variability of these estimates: a standard error of 0 for serotonin indicated no variability in the estimate, while a standard error of 0.067 for CDR-SB indicated some level of variability in the estimate for that variable. The 95% confidence interval for CDR-SB ranged from -0.322 to -0.073, reinforcing the negative correlation. Overall, these results highlighted a positive significant relationship between serotonin levels and cognitive function, suggesting serotonin’s potential role in cognitive functions. (Table 3)

**Table 3:** Correlations between serotonin level and cognitive functions measured by CDR-SB.

| Variable | Serotonin | CDR |
| --- | --- | --- |
| Pearson Correlation | 1 | **-0.198\*** |
| Significance (2-tailed) |  | 0.023 |
| Sum of Squares and Cross-products | 21.593 | -25.287 |
| Covariance | 0.164 | -0.192 |
| Bootstrap Bias | 0 | -0.004 |
| Standard Error | 0 | 0.067 |
| BCa 95% Confidence Interval |  |  |
| Lower |  | -0.322 |
| Upper |  | -0.073 |

\* Correlation is significant at the 0.05 level (2-tailed).

### 3.3. Total Brain Volume and Bilateral Hippocampal Volumes Correlated with Cognitive Functions Measured by CDR-SB

The findings revealed correlations between the CDR-SB score, total brain volume, and VBM measures of the left and right hippocampus in participants. Higher CDR scores, indicating greater cognitive impairment, were significantly correlated with reductions in total brain volume (r = -0.223, p = 0.010), left hippocampal volume (r = -0.246, p = 0.004), and right hippocampal volume (r = -0.308, p < 0.001), with a stronger relationship observed in the right hippocampus. Bootstrap analysis, used for verification, showed minimal bias and confirmed the significance of these correlations. (Table 4)

**Table 4:** Correlations between various brain volume measurements and cognitive functions measured by CDR-SB.

| Variables | CDR | Bias | Std. Error | BCa 95% Confidence Interval (Lower) | BCa 95% Confidence Interval (Upper) |
| --- | --- | --- | --- | --- | --- |
| CDR | 1 | 0 | 0 | - | - |
| Total Brain Volume | **-0.223\*\*** ($p = 0.010$) | 0.001 | 0.080 | -0.365 | -0.063 |
| Left Hippocampus | **-0.246\*\*** ($p = 0.004$) | 0.004 | 0.081 | -0.392 | -0.070 |
| Right Hippocampus | **-0.308\*\*** ($p < 0.001$) | 0.003 | 0.085 | -0.470 | -0.134 |

\*\*Correlation is significant at the 0.01 level (2-tailed).

### 3.4. Serotonin Level Correlated with Various Neurotrophic and Nerve Growth Factors Involved in Neurogenesis and Neuroplasticity

The findings showed that serotonin has significant positive correlations with BMP-6 (r = 0.173, p = 0.047), CNTF (r = 0.216, p = 0.013), FGF-4 (r = 0.176, p = 0.043), and MMP-1 (r = 0.202, p = 0.019), suggesting that higher serotonin levels were associated with increased levels of these factors. BMP-6 strongly correlated with CNTF (r = 0.750, p < 0.001), FGF-4 (r = 0.734, p < 0.001), and MMP-1 (r = 0.446, p < 0.001), indicating significant co-regulation. Besides, CNTF and FGF-4 were strongly linked (r = 0.714, p < 0.001), and both showed positive correlations with MMP-1. The bootstrap analysis validated these findings with minimal biases and standard errors, providing confidence in the stability of these relationships. (Table 5) (Figure 1)

**Table 5:** Correlations between serotonin level and various neurotrophic and nerve growth factors.

| Variables | Pearson Correlation (r) | p-value | Bias | Std. Error | BCa 95% Confidence Interval (Lower) | BCa 95% Confidence Interval (Upper) |
| --- | --- | --- | --- | --- | --- | --- |
| **BMP-6** | **0.173\*** | 0.047 | -0.013 | 0.126 | -0.040 | 0.370 |
| **CNTF** | **0.216\*** | 0.013 | -0.007 | 0.106 | 0.018 | 0.399 |
| **FGF-4** | **0.176\*** | 0.043 | -0.007 | 0.108 | -0.040 | 0.361 |
| **MMP-1** | **0.202\*** | 0.019 | -0.013 | 0.131 | -0.046 | 0.428 |

\*Correlation is significant at the 0.05 level (2-tailed); BMP-6: Bone Morphogenetic Protein 6; CNTF: Ciliary Neurotrophic Factor; FGF-4: Fibroblast Growth Factor 4; MMP-1: Matrix Metalloproteinase-1.

**Figure 1.**
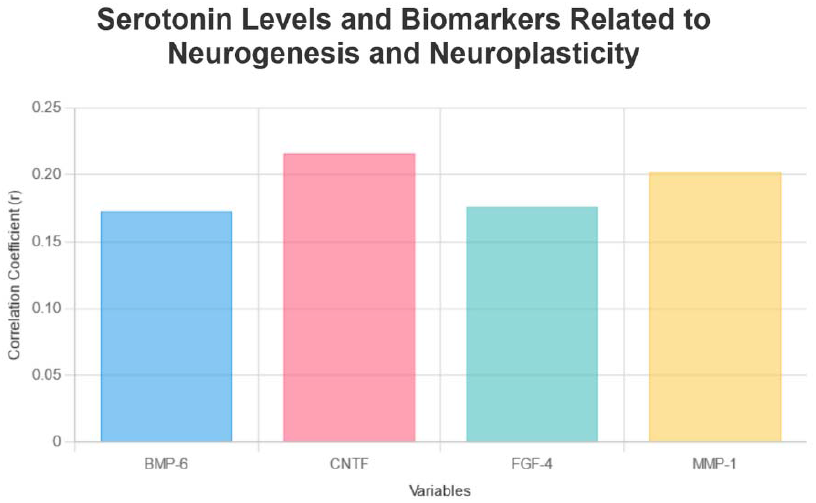
The bar chart presents the correlation coefficients of key biomarkers—BMP-6 (0.173), CNTF (0.216), FGF-4 (0.176), and MMP-1 (0.202)—associated with serotonin levels. These biomarkers play significant roles in neurogenesis and neuroplasticity, indicating their potential impact on brain health and development.

## 4. Discussion

The study on AD patients uncovered several significant relationships that emphasize the role of serotonin in brain health and cognitive function. A key finding, for the first time, was the positive correlation between serotonin levels and various biomarkers and neurotrophic factors, including CNTF, FGF-4, BMP-6, and MMP-1. These factors are involved in neurogenesis and neuroplasticity, suggesting that higher serotonin levels may stimulate their production, thereby promoting brain resilience and supporting cognitive health.

In addition, the research revealed a significant negative correlation between serotonin levels and cognitive impairment, as measured by the CDR-SB. Higher serotonin levels were associated with better cognitive function, suggesting a protective role against cognitive decline in AD and emphasizing its importance in cognitive health.

The study also found a positive association between serotonin levels and brain volume. This means that participants with higher serotonin levels exhibited increased total brain volume and larger hippocampal volumes, suggesting that elevated serotonin may help preserve or enhance brain structure. Moreover, the study demonstrated that reductions in brain volume, particularly in the hippocampus, were linked to greater cognitive impairment, further reinforcing the idea that maintaining brain and hippocampal size is crucial for cognitive preservation in AD patients.

Overall, the findings emphasized the interconnected roles of serotonin, brain structure, cognitive function, biomarkers, and neurotrophic factors in AD. They also suggested potential therapeutic strategies that focus on boosting serotonin levels to support brain health and enhance cognitive performance.

The findings of this study are consistent with earlier research emphasizing serotonin’s vital role in maintaining brain health and cognitive function. Both this research and prior studies demonstrated a clear link between higher serotonin levels and better cognitive performance, suggesting that serotonin plays a crucial role in enhancing cognitive outcomes. This connection highlights serotonin’s significance beyond its traditional function as a neurotransmitter, positioning it as a central component in the neurobiological mechanisms that support cognitive health [42-47].

Serotonin plays a crucial role in both physiological and cognitive functions by modulating neural activity through its widespread receptor subtypes. It influences key brain regions like the hippocampus and prefrontal cortex, which are essential for learning, memory, and decision-making [48-52]. The serotonergic system interacts with other neurotransmitters, such as dopamine, acetylcholine, and glutamate, contributing to cognitive processes [52]. In disorders like schizophrenia [53-57] and AD [58-63], disrupted serotonergic pathways were linked to cognitive deficits, including impaired memory and reasoning. Thus, therapeutic strategies targeting serotonin receptors, such as 5-HTR3 and 5-HTR4, showed potential in improving cognition by regulating neurotransmitter release and reducing pathological markers like amyloid-β in AD [63-66].

The current findings expand previous knowledge by highlighting serotonin-linked biomarkers—CNTF, FGF-4, BMP-6, and MMP-1—that have been largely underexplored in earlier studies. These biomarkers are crucial for neurogenesis and synaptic plasticity [67-73], shedding light on potential mechanisms through which serotonin may influence cognitive health and brain resilience. However, the precise pathways by which serotonin increases these biomarkers remain unclear. While there is evidence that serotonin affects the expression of neurotrophic factors like BDNF [74, 75], its interactions with CNTF, FGF-4, BMP-6, and MMP-1 are not yet fully understood.

CNTF is a neurotrophic factor crucial for neuron survival and differentiation, primarily activating the JAK/STAT pathway [76-78] to maintain neuronal health and induce neurogenesis, particularly in conditions like AD [79-81]. FGF-4 similarly promotes the proliferation and differentiation of neural progenitor cells through FGF receptors and the MAPK pathway, supporting neurogenesis and cognitive resilience [82-85]. BMP-6 may contribute to neuronal development and synaptic remodeling via Smad signaling, enhancing cognitive adaptability [86-89]. MMP-1 plays also a key role in extracellular matrix remodeling, enabling structural changes in synapses critical for synaptic plasticity [90, 91]. In short, serotonin’s influence on CNTF, FGF-4, BMP-6, and MMP-1 supports neurogenesis, neuronal survival, and synaptic plasticity, fostering cognitive resilience. However, the precise mechanisms by which serotonin mediates these effects remain unclear, requiring further research to fully understand its role in neurodegenerative diseases.

It is of note that some former findings highlighted the increased expression of BMP6 in the hippocampus of both human AD patients and APP transgenic mice, indicating its potential role in disrupting neurogenesis. For instance, qRT-PCR analysis revealed elevated BMP6 mRNA levels, which were corroborated by immunoblotting and immunohistochemical methods. The accumulation of BMP6 protein around amyloid plaques links its overexpression to amyloid-β pathology. In vitro studies further demonstrated that Aβ elevates BMP6 expression, subsequently inhibiting neural progenitor cell proliferation without inducing toxicity. These results suggest that Aβ-driven BMP6 upregulation may impair hippocampal neurogenesis, contributing to AD progression [92, 93]. Normalizing BMP6 levels could present a therapeutic strategy to mitigate neurogenic deficits in AD [93]. Similarly, another study found that increased BMP signaling in aged mice inhibits neural progenitor cell proliferation, but reducing BMP signaling restores neurogenesis, highlighting potential targets for treating age-related cognitive decline [94]. However, the significance of neurogenesis in cognitive function, particularly in the aging brain, remains unclear, as many AD-affected regions, like the neocortex, are non-neurogenic; thus, further investigation is needed to determine how such biomarkers can promote neurogenesis and potentially benefit AD patients.

The study found that higher serotonin levels in AD patients were positively associated with increased total brain and hippocampal volumes, indicating serotonin’s role in preserving brain structure. This aligns with earlier studies, though conducted on healthy individuals or other diseases, that emphasized serotonin’s influence on GMV and hippocampal size. Specifically, serotonin receptor binding, particularly 5-HT1A, was positively correlated with GMV, indicating its role in maintaining brain integrity [28]. In psychiatric conditions, altered serotonin receptor densities were linked to reduced GMV, impaired neurogenesis, and cognitive deficits [29, 30]. Genetic factors, like the His452Tyr variant in the 5-HT2A receptor, were also linked to reduced hippocampal volume and poorer memory performance [31]. However, no such correlation was found in individuals with autism spectrum disorder, highlighting variability in serotonin’s effects across conditions [32]. These findings underscore serotonin’s complex role in brain structure and cognitive function, calling for further research.

The study had several strengths that enhanced our understanding of serotonin’s role in AD. One key strength was its focus on specific brain regions, particularly the hippocampus, which is crucial in AD pathology. This targeted approach added significant relevance to the findings. Besides, the study explored the connections between serotonin levels and neurotrophic factors like BMP-6, CNTF, and FGF-4. This study added a valuable layer of molecular insight, suggesting that serotonin might have influenced neurogenesis and neuroplasticity in the context of neurodegeneration. The use of bootstrap analysis also increased the reliability of the results, providing more robust estimates of the relationships among the variables studied.

However, there were notable limitations in this research that needed to be considered. The cross-sectional design made it difficult to establish causal relationships between serotonin levels and brain structure or cognitive function, limiting the ability to determine the direction of these associations. Further, the study had a relatively small sample size, which raised concerns about the statistical power of the findings. The sample’s demographic composition did not fully represent the broader AD population, particularly regarding gender balance and age distribution, which could have limited the applicability of the results. Moreover, the presence of non-normal distributions in variables like CDR-SB, ADAS-Cog, and MMSE could have impacted the accuracy of parametric statistical tests, potentially leading to biased conclusions. Lastly, the study did not include genetic analyses, leaving out the possibility of genetic factors influencing serotonin’s effects on brain structure and function. Future studies should address these limitations by adopting longitudinal designs, increasing sample size, ensuring demographic diversity, applying appropriate statistical methods, and incorporating genetic analyses to achieve more convergent findings.

## 5. Conclusion

The findings revealed that serotonin plays a crucial role in influencing brain structure and cognitive functions in individuals with AD. Specifically, higher serotonin levels were associated with larger brain and hippocampal volumes, which in turn correlated with better cognitive performance. Furthermore, the positive relationships between serotonin and biomarkers, related to neurogenesis and neuroplasticity, such as CNTF, FGF-4, BMP-6, and MMP-1, indicated that serotonin may enhance neurogenesis and neuroplasticity, providing potential therapeutic avenues for improving cognitive function in AD. These insights underscore the importance of serotonin in the context of neurodegeneration and highlight areas for further research into its therapeutic potential.

## Abbreviations

AD: Alzheimer’s disease
DG: dentate gyrus
SVZ: subventricular zone
BDNF: brain-derived neurotrophic factor
CNTF: ciliary neurotrophic factor
FGF-4: fibroblast growth factor 4
BMP-6: bone morphogenetic protein 6
MMP-1: matrix metalloproteinase-1
GMV: gray matter volume
CDR: clinical dementia rating
ADAS-Cog: Alzheimer’s disease assessment scale-cognitive subscale
MMSE: mini-mental state examination
QC: quality control
VBM: voxel-based morphometry
CSF: cerebrospinal fluid

## Declaration sections

## Acknowledgments

Data collection and sharing for this project was funded by the Alzheimer’s Disease Neuroimaging Initiative (ADNI) (National Institutes of Health Grant U01 AG024904) and DOD ADNI (Department of Defense award number W81XWH-12-2-0012). ADNI is funded by the National Institute on Aging, the National Institute of Biomedical Imaging and Bioengineering, and through generous contributions from the following: AbbVie, Alzheimer’s Association; Alzheimer’s Drug Discovery Foundation; Araclon Biotech; BioClinica, Inc.; Biogen; Bristol-Myers Squibb Company; CereSpir, Inc.; Cogstate; Eisai Inc.; Elan Pharmaceuticals, Inc.; Eli Lilly and Company; EuroImmun; F. Hoffmann-La Roche Ltd and its affiliated company Genentech, Inc.; Fujirebio; GE Healthcare; IXICO Ltd.; Janssen Alzheimer Immunotherapy Research & Development, LLC.; Johnson & Johnson Pharmaceutical Research & Development LLC.; Lumosity; Lundbeck; Merck & Co., Inc.; Meso Scale Diagnostics, LLC.; NeuroRx Research; Neurotrack Technologies; Novartis Pharmaceuticals Corporation; Pfizer Inc.; Piramal Imaging; Servier; Takeda Pharmaceutical Company; and Transition Therapeutics. The Canadian Institutes of Health Research is providing funds to support ADNI clinical sites in Canada. Private sector contributions are facilitated by the Foundation for the National Institutes of Health (www.fnih.org). The grantee organization is the Northern California Institute for Research and Education, and the study is coordinated by the Alzheimer’s Therapeutic Research Institute at the University of Southern California. ADNI data are disseminated by the Laboratory for Neuro Imaging at the University of Southern California.

## Consent to Participate

This study was conducted using ADNI data. The ADNI study is ethically approved and operated in accordance with the Declaration of Helsinki, 1964.

## Consent for Publication

Not applicable.

## Authors’ Contributions

The entire research, including all its parts, was conducted by A.A.

## Availability of Data and Materials

The data used in this research was obtained from Alzheimer’s Disease Neuroimaging Initiative (ADNI) and is available with permission to all researchers.

